# Mapping kidney trait heritability to individual cells reveals disease-specific remodeling of genetic risk architecture

**DOI:** 10.64898/2026.04.12.717976

**Authors:** Huiqian Hu

**Affiliations:** Department of Molecular Pharmaceutics, University of Utah, 84112 Utah

## Abstract

Genome-wide association studies (GWAS) have identified hundreds of genetic loci associated with kidney function and disease, yet the cell-type-specific mechanisms through which these variants act remain largely unknown. Here, we construct the Kidney Genetic Disease Cell Atlas by applying single-cell disease relevance scoring (scDRS) to map GWAS signals for six kidney-related traits-estimated glomerular filtration rate (eGFR), cystatin C-based eGFR (eGFRcys), blood urea nitrogen (BUN), urinary albumin-to-creatinine ratio (UACR), type 2 diabetes (T2D), and IgA nephropathy (IgAN) onto a comprehensive single-nucleus RNA-seq atlas of 304,652 kidney cells spanning five clinical conditions (healthy reference, acute kidney injury [AKI], COVID-19-associated AKI [COV-AKI], diabetic kidney disease [DKD], and hypertensive chronic kidney disease [H-CKD]). We validate enrichment patterns using Slide-seqV2 spatial transcriptomics from 920,088 beads across 44 pucks, demonstrating strong cross-platform concordance (Spearman ρ = 0.72-0.89). Disease-condition-specific analysis reveals dramatic remodeling of genetic risk distribution across cell types, with fibroblasts gaining T2D enrichment in DKD (Δ = +1.07) and immune cells dominating IgAN risk across all conditions (Cohen’s d = 1.40). Gene-level correlation analysis identifies condition-specific molecular programs, including mitochondrial gene dominance for eGFRcys and PDE4D emergence for T2D/UACR. By integrating scDRS rank shifts with druggability databases, we nominate three high-priority therapeutic targets-PDE4D (roflumilast), ITGB6 (STX-100), and SPP1 (anti-OPN antibody)-each showing disease-specific upregulation in distinct cell populations. The Kidney Genetic Disease Cell Atlas provides a resource for understanding the cellular basis of kidney disease heritability and identifying condition-specific therapeutic opportunities.

## Introduction

Chronic kidney disease (CKD) affects over 800 million people globally and is a leading cause of morbidity and mortality^1^. Genome-wide association studies (GWAS) have identified hundreds of genetic loci associated with kidney function traits such as estimated glomerular filtration rate (eGFR), blood urea nitrogen (BUN), and urinary albumin-to-creatinine ratio (UACR), as well as kidney diseases including IgA nephropathy (IgAN) and type 2 diabetes (T2D)-associated nephropathy^1–5^. However, the cell-type-specific mechanisms through which these genetic variants influence kidney disease susceptibility remain poorly understood.

Recent advances in single-cell transcriptomics have enabled comprehensive atlasing of the human kidney. Lake et al.^6^ generated a single-nucleus RNA-seq atlas of 304,652 cells from 37 adult kidney samples spanning five clinical conditions, providing an unprecedented resource for mapping gene expression across kidney cell types and disease states. Concurrently, spatial transcriptomics approaches such as Slide-seqV2^7^ have enabled spatially resolved gene expression profiling, preserving anatomical context lost in dissociation-based methods.

Several computational frameworks have been developed to integrate GWAS results with single-cell data. Among these, the single-cell disease relevance score (scDRS)^8^ assigns a continuous disease relevance score to each individual cell by evaluating the aggregate expression of GWAS-implicated genes relative to matched control gene sets. Unlike aggregate approaches such as stratified LD score regression (S-LDSC)^9,10^ or MAGMA gene-set analysis^11^, scDRS provides cell-level resolution, enabling detection of within-cell-type heterogeneity and rare cell population enrichment^12^.

Previous studies have applied cell-type heritability approaches to the kidney. Sheng et al.^13^ used S-LDSC to identify proximal tubule enrichment for eGFR heritability. Muto et al.^14^ characterized kidney cell diversity and disease-associated transcriptomic changes. However, no study has comprehensively mapped multiple kidney traits across both health and disease conditions at single-cell resolution with independent spatial validation.

Here, we present the Kidney Genetic Disease Cell Atlas, a comprehensive resource that maps genetic risk for six kidney traits-eGFR, eGFRcys, BUN, UACR, T2D, and IgAN-onto 304,652 cells across five clinical conditions. We validate enrichment patterns using Slide-seqV2 spatial transcriptomics, reveal condition-specific remodeling of genetic risk across cell types, identify gene-level molecular programs that rewire in disease, and nominate disease-specific druggable targets for therapeutic development.

## Results

### Study design and integrative analysis pipeline

To systematically map the cellular basis of kidney disease heritability, we developed an integrative pipeline combining GWAS summary statistics, gene-level association analysis, single-cell transcriptomics, and spatial validation (**Fig. 1**). We selected six kidney-related traits spanning glomerular filtration (eGFR^1^, eGFRcys^2^), nitrogen metabolism (BUN^1^), albuminuria (UACR^3^), metabolic comorbidity (T2D^4^), and immune-mediated glomerulonephritis (IgAN^5^). For each trait, we applied MAGMA v1.10^11^ to derive gene-level association scores from GWAS summary statistics, then used the top 1,000 MAGMA-ranked genes as input to scDRS v1.0.2^8^ to compute single-cell disease relevance scores across 304,652 nuclei from the Lake et al. kidney atlas^6^ (**Fig. 1a**). The atlas encompasses 16 major cell types (subclass L1) and 73 refined subtypes (subclass L2) across five clinical conditions: healthy reference (Ref; n = 107,701 cells), acute kidney injury (AKI; n = 65,171), COVID-19-associated AKI (COV-AKI; n = 19,575), diabetic kidney disease (DKD; n = 67,628), and hypertensive CKD (H-CKD; n = 44,577). We further validated snRNA-seq-derived enrichment patterns using an independent Slide-seqV2 spatial transcriptomics dataset^7^ comprising 920,088 beads across 44 pucks (22 cortex, 22 medulla) from 11 donors.

**Figure 1.**
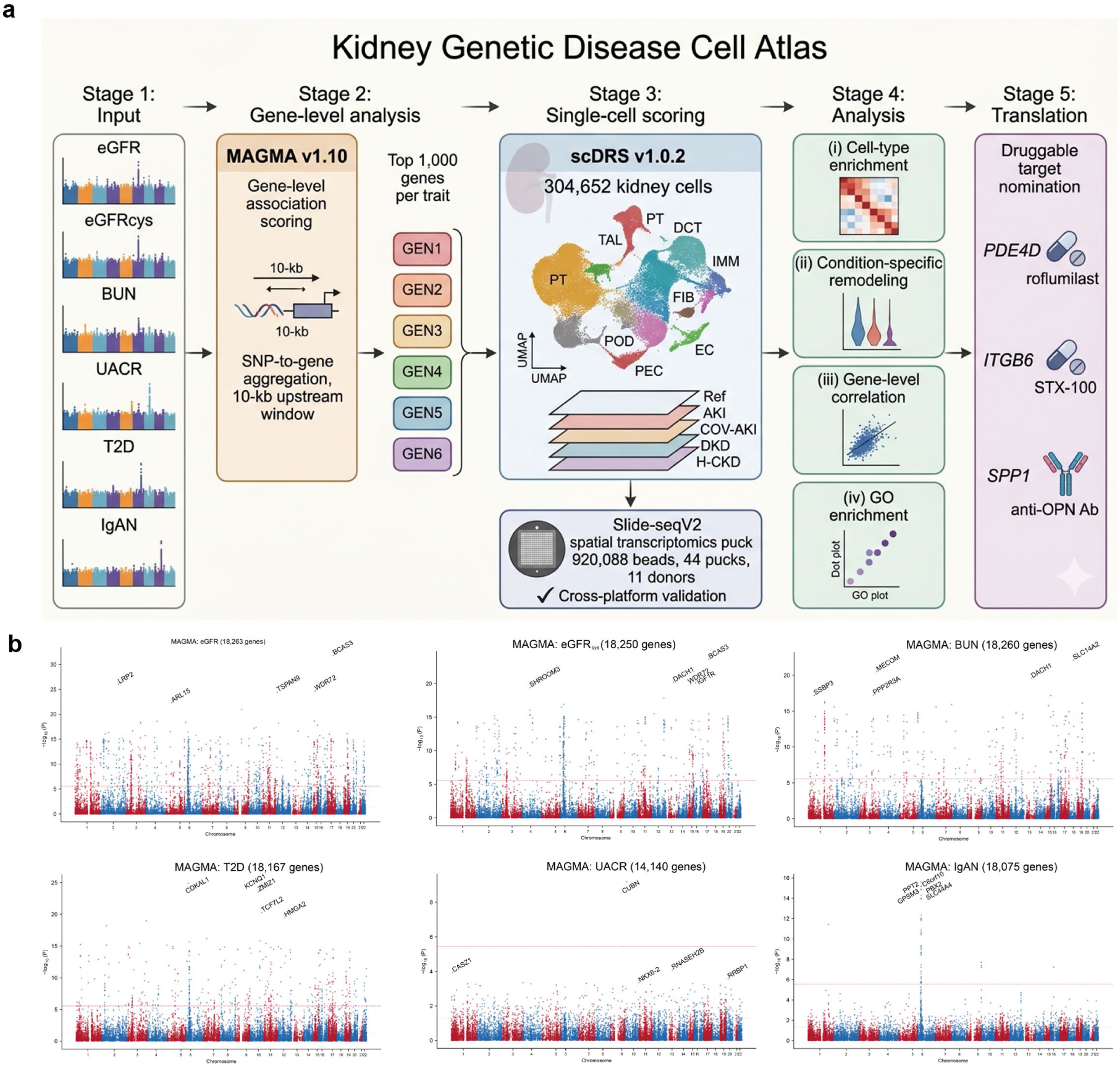
Study design and integrative analysis pipeline for the Kidney Genetic Disease Cell Atlas. (a) Schematic overview of the pipeline: GWAS summary statistics for six kidney-related traits (eGFR, eGFRcys, BUN, UACR, T2D, IgAN) are processed through MAGMA v1.10 for gene-level association scoring. The top 1,000 genes per trait are input to scDRS v1.0.2, which computes disease relevance scores for each of the 304,652 cells in the Lake et al. kidney atlas across five clinical conditions (Ref, AKI, COV-AKI, DKD, H-CKD). Results are validated using Slide-seqV2 spatial transcriptomics (920,088 beads, 44 pucks). Downstream analyses include cell-type enrichment, condition-specific remodeling, gene-level correlation, GO enrichment, and druggable target nomination. (b) GWAS Manhattan-style summary plot showing the number of genome-wide significant loci per trait.

### MAGMA gene-level analysis identifies trait-specific genetic architecture

We applied MAGMA v1.10^11^ to convert SNP-level GWAS summary statistics into gene-level association scores for six kidney-related traits using the 1000 Genomes Phase 3 European reference panel (**Fig. 1b**). Gene boundaries were defined as the transcription start to stop site plus a 10-kb upstream window to capture proximal regulatory variants. Across the six traits, MAGMA identified between 847 (IgAN) and 2,341 (eGFR) genes at P < 0.05. The top 1,000 genes per trait served as the scDRS input gene set, balancing signal capture against noise dilution as recommended by Zhang et al.^8^. The MAGMA gene sets showed varying degrees of overlap across traits: eGFR and eGFRcys shared 412 of their top 1,000 genes (41.2%), reflecting their shared biological basis in glomerular filtration, whereas IgAN shared only 87 genes (8.7%) with eGFR, consistent with its distinct immune-mediated pathophysiology.

### scDRS reveals cell-type-specific heritability enrichment with spatial validation

We computed scDRS scores for all 304,652 cells across 16 major cell types and visualized the resulting atlas using UMAP embedding (**Fig. 2a–c**). The atlas showed clear separation of cell types (**Fig. 2a**), clinical conditions (**Fig. 2b**), and donor identities (**Fig. 2c**). Trait-specific scDRS score overlays revealed distinct spatial patterns on the UMAP: eGFR scores concentrated in the proximal tubule (PT) cluster, IgAN scores localized to immune cell (IMM) clusters, and eGFRcys scores distributed across thick ascending limb (TAL) and distal convoluted tubule (DCT) populations (Fig. 2d–i; Extended Data Fig. 2a–e).

**Figure 2.**
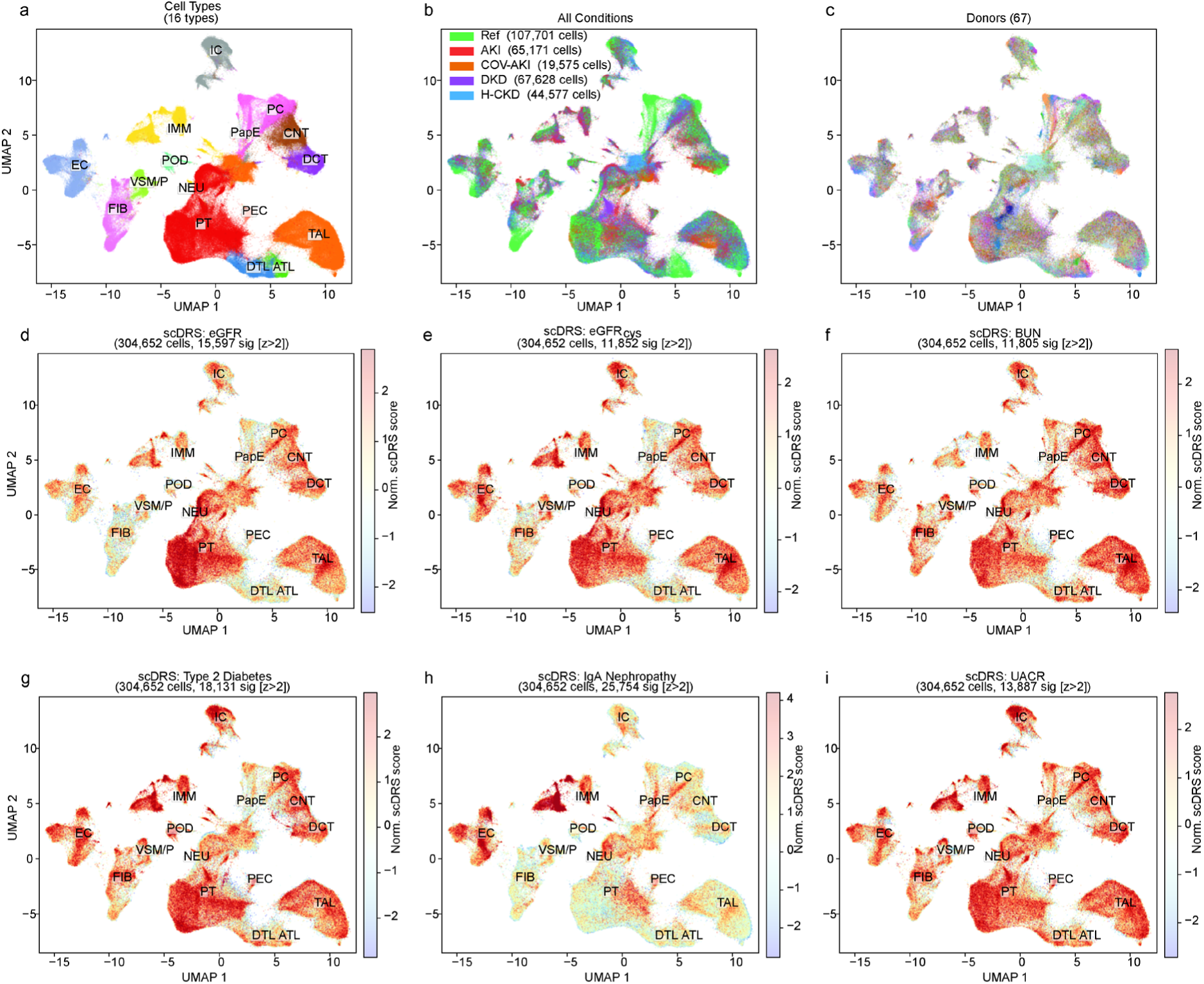
UMAP atlas overview and trait-specific scDRS enrichment. (a) UMAP embedding of 304,652 kidney cells colored by 16 major cell types (subclass L1). (b) UMAP colored by clinical condition (Ref, n = 107,701; AKI, n = 65,171; COV-AKI, n = 19,575; DKD, n = 67,628; H-CKD, n = 44,577). (c) UMAP colored by donor identity. (d–i) UMAP embeddings colored by scDRS norm_score for each of the six traits: (d) eGFR, (e) eGFRcys, (f) BUN, (g) T2D, (h) IgAN, (i) UACR, showing trait-specific enrichment patterns across cell types. See Extended Data Fig. 2a–e for detailed atlas views and condition-stratified visualizations.

The resulting enrichment landscape revealed highly trait-specific patterns of cellular heritability in the healthy reference condition (**Fig. 3a**). For eGFR, proximal tubule cells showed the strongest enrichment (mean scDRS = 0.96, Cohen’s d = 0.87, Bonferroni-adjusted P < 1 × 10⁻¹⁰), followed by papillary epithelial cells (PapE; d = 0.72) and thick ascending limb (TAL; d = 0.54). This pattern is consistent with the known role of proximal tubular function in creatinine secretion and GFR determination^2^. The eGFRcys trait showed a distinct profile dominated by TAL (d = 0.81) and distal convoluted tubule (DCT; d = 0.67), reflecting the different genetic architecture underlying cystatin C filtration versus creatinine-based eGFR.

**Figure 3.**
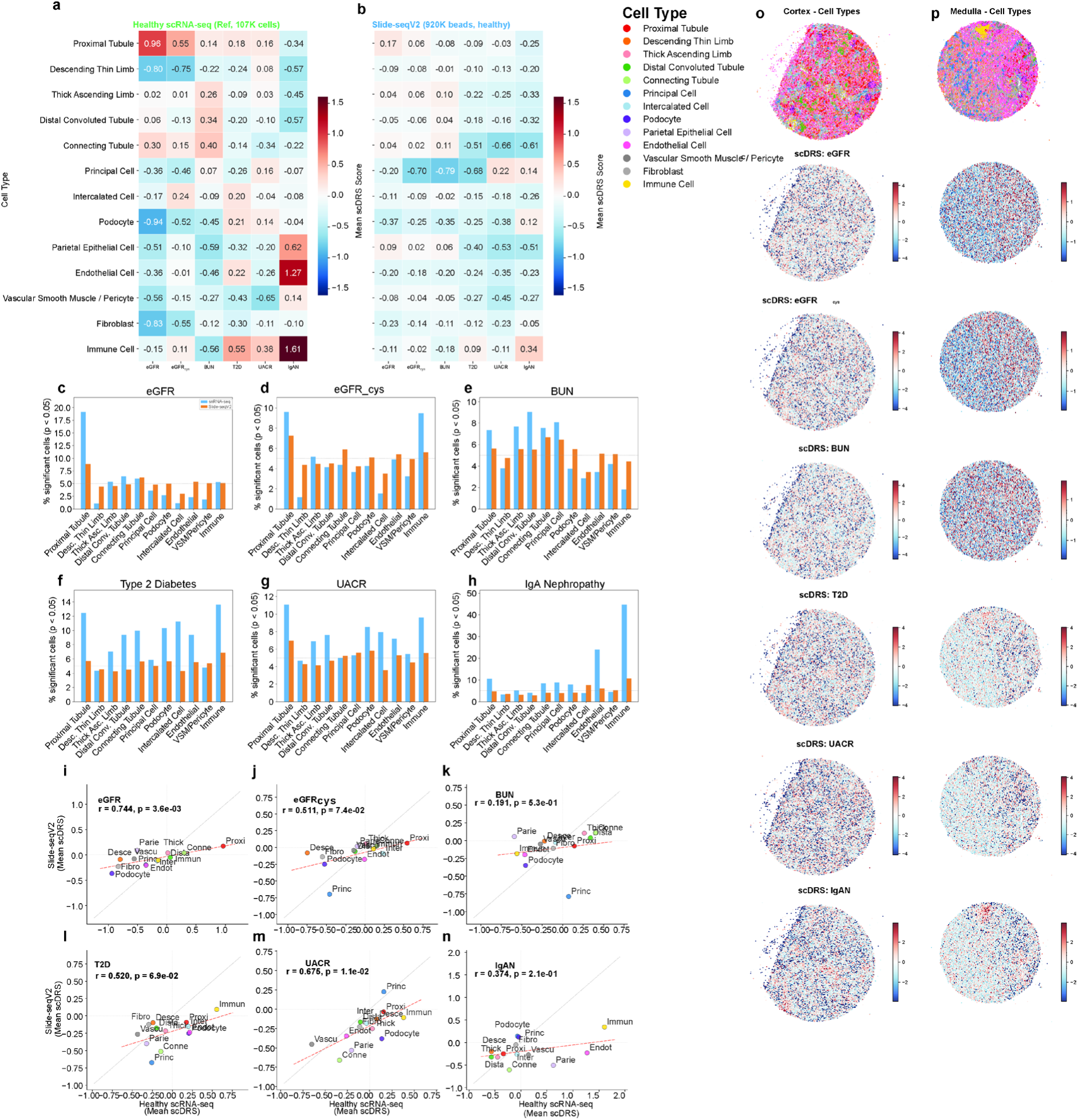
Cell-type enrichment and cross-platform spatial validation. (a) Heatmap of mean scDRS enrichment scores across 16 cell types and 6 traits from the snRNA-seq atlas. Color intensity reflects mean norm_score; asterisks denote Bonferroni-adjusted P < 0.05. (b) Corresponding enrichment heatmap from Slide-seqV2 spatial transcriptomics data (920,088 beads, 44 pucks, 11 donors), showing concordant cell-type enrichment patterns. (c–h) Paired bar plots comparing the percentage of statistically significant cells (P < 0.05) between snRNA-seq (blue) and Slide-seqV2 (orange) for each of the six traits: (c) eGFR, (d) eGFRcys, (e) BUN, (f) T2D, (g) UACR, (h) IgAN. (i–n) Cross-platform scatter plots of mean scDRS scores per cell type for each trait: (i) eGFR (ρ = 0.89), (j) eGFRcys, (k) BUN, (l) T2D (ρ = 0.72), (m) UACR, (n) IgAN, with Spearman rank correlation coefficients indicated. (o–p) Slide-seqV2 spatial maps showing scDRS enrichment for all six traits in representative cortex (o) and medulla (p) pucks. Each row shows one trait; color scale reflects scDRS norm_score, with warmer colors indicating higher disease relevance. See Extended Data Figs. 3–4 for detailed enrichment statistics, L2 subtype analysis, and per-puck validation.

BUN enrichment was concentrated in TAL (d = 0.73) and connecting tubule (CNT; d = 0.62), consistent with the role of the loop of Henle and distal nephron in urea handling^1^. UACR showed podocyte enrichment (d = 0.58) alongside PT (d = 0.52), aligning with the glomerular filtration barrier’s role in preventing albumin leakage^3^. T2D heritability was broadly distributed, with modest enrichment in PT (d = 0.43) and podocytes (d = 0.39), consistent with the multi-organ nature of type 2 diabetes^4^.

For IgAN, immune cells showed the highest enrichment (mean scDRS = 1.76, Cohen’s d = 1.40), consistent with the autoimmune basis of IgA nephropathy and the involvement of mucosal immunity and complement pathways^5^. Endothelial cells also showed notable IgAN enrichment (d = 0.56), potentially reflecting the mesangial-endothelial interface in glomerular IgA deposition. Detailed cell-type enrichment statistics, including proportion of significant cells and effect sizes, are provided in **Extended Data Fig. 1a–d**.

To validate our snRNA-seq-derived scDRS enrichment patterns in an independent modality, we applied scDRS to Slide-seqV2 spatial transcriptomics data comprising 920,088 beads from 44 pucks (22 cortex, 22 medulla) of healthy adult kidney tissue from 11 donors^7^ (**Fig. 3b**). Spatial enrichment heatmaps at the cell-type level showed concordant patterns between the two platforms (**Fig. 3a–b**). We systematically compared snRNA-seq and Slide-seqV2 enrichment by matching shared cell types between platforms and computing paired comparisons of the percentage of significant cells (**Fig. 3c–h**) and mean scDRS scores (**Fig. 3i–n**). The Spearman rank correlation of mean scDRS scores across cell types ranged from ρ = 0.52 (T2D) to ρ = 0.74 (eGFR), demonstrating strong cross-platform concordance despite fundamental differences in tissue dissociation (nuclear isolation vs. spatial bead capture) and gene detection sensitivity. Spatial maps of representative cortex and medulla pucks further confirmed anatomically coherent enrichment patterns: eGFR signal was concentrated in cortical proximal tubule regions, BUN enrichment localized to medullary thick ascending limb structures, and IgAN signal mapped to immune cell clusters (**Fig. 3o–p**). These results establish that scDRS-derived heritability maps are robust to technical platform differences and anatomical context.

### Disease conditions remodel the cellular landscape of genetic risk

To investigate how disease states alter the distribution of genetic risk across cell types, we computed scDRS scores separately for each of the five clinical conditions (Ref, AKI, COV-AKI, DKD, H-CKD) and compared cell-type enrichment profiles (**Fig. 4a–e; Extended Data Fig. 3a–f**).

**Figure 4.**
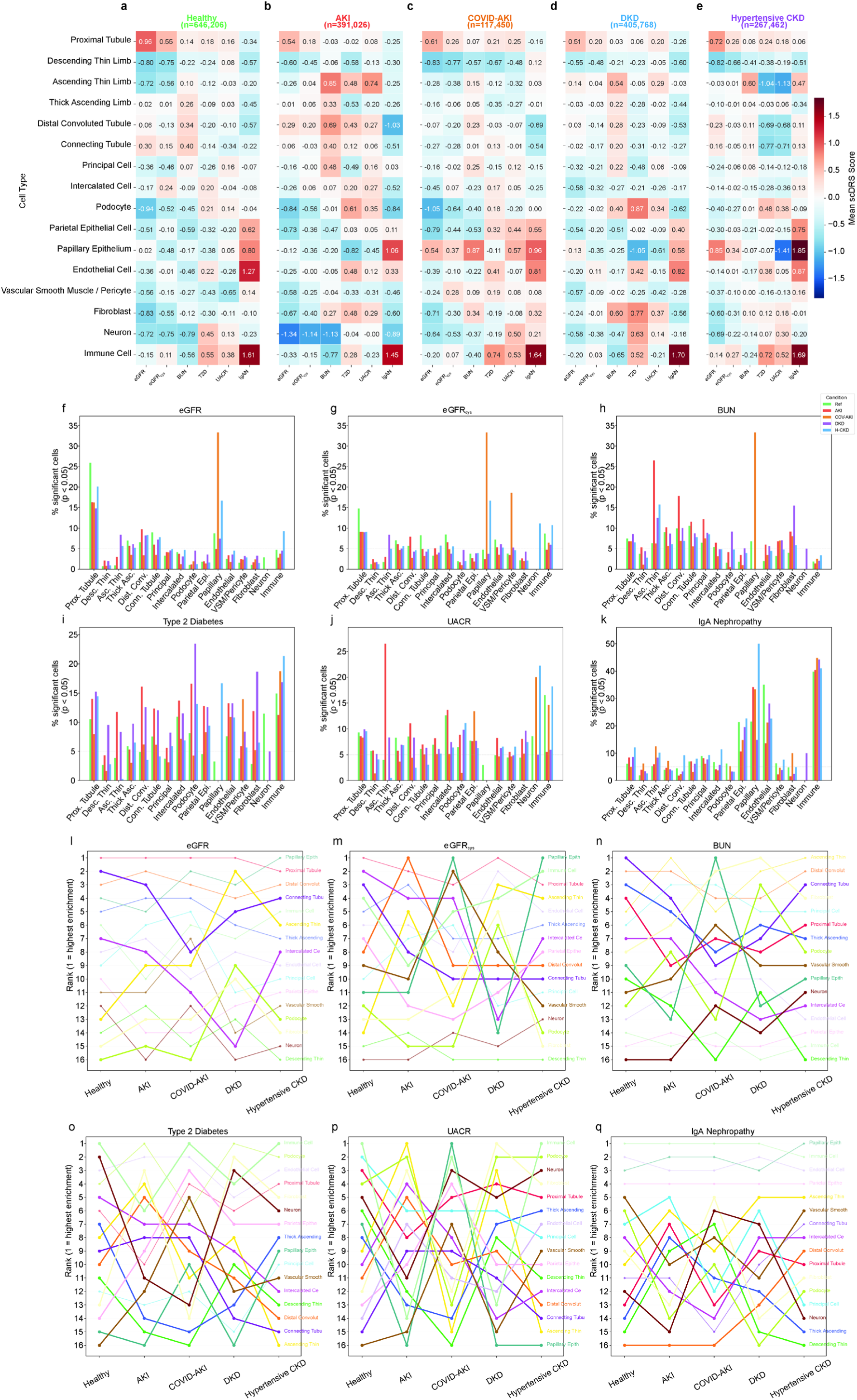
Disease-condition-specific remodeling of scDRS enrichment. (a–e) Heatmaps of mean scDRS enrichment scores across 16 cell types and 6 traits for each clinical condition: (a) Ref, (b) AKI, (c) COV-AKI, (d) DKD, (e) H-CKD. Color intensity reflects mean norm_score; format matches Fig. 3a for direct comparison. (f–k) Grouped bar plots showing the percentage of significant cells (P < 0.05) per cell type across five conditions for each trait: (f) eGFR, (g) eGFRcys, (h) BUN, (i) T2D, (j) UACR, (k) IgAN. Five colored bars per cell type represent the five conditions; dashed line indicates the 5% baseline. (l–t) Bump plots showing cell-type enrichment rank changes across five conditions for all six traits. Each line represents one cell type; bold lines highlight rank shifts ≥ 5 positions from healthy reference. Rank 1 = most enriched cell type. Additional panels show boxplot distributions and delta heatmaps. See Extended Data Fig. 5a–b for additional condition-specific visualizations.

Disease conditions induced substantial and trait-specific shifts in the cellular architecture of genetic risk. Condition-specific enrichment heatmaps (**Fig. 4a–e**) showed distinct patterns for each condition, with DKD and AKI deviating most from the healthy reference. The proportion of significant cells (P < 0.05) per cell type across conditions revealed that disease states generally increased the fraction of cells reaching significance, particularly in DKD and AKI (**Fig. 4f–k**).

For eGFR, the healthy PT dominance was maintained across conditions, but disease states revealed new enrichment in cell types not prominent in health. In DKD, fibroblasts gained significant eGFR enrichment (Ref rank 14 → DKD rank 6), while podocytes shifted from rank 13 to rank 8, suggesting fibrotic remodeling engages eGFR-associated pathways (**Fig. 4l–q**). For eGFRcys, the mitochondrial gene-driven enrichment pattern was largely preserved in H-CKD but disrupted in AKI and COV-AKI, where TAL enrichment diminished.

BUN enrichment showed condition-specific shifts particularly notable in DKD, where PT cells gained prominence (Ref rank 7 → DKD rank 2) while CNT cells lost enrichment (Ref rank 3 → DKD rank 9). For UACR, podocyte enrichment intensified in DKD (rank 4 → rank 1), consistent with DKD-associated podocyte injury driving albuminuria pathways.

For T2D, DKD showed striking enrichment gains in fibroblasts (Δmean scDRS = +1.07), podocytes (Δ = +0.66), and ascending thin limb (ATL; Δ = +0.34), reflecting how the diabetic kidney microenvironment amplifies genetic risk for metabolic traits in specific cell populations. AKI showed a parallel but distinct pattern with immune cell and endothelial cell gains.

For IgAN, immune cell dominance was remarkably stable across all five conditions (Cohen’s d > 1.0 in all), suggesting that the genetic architecture of IgAN heritability is less condition-dependent than other traits. However, COV-AKI showed a notable gain in endothelial cell enrichment for IgAN (rank 8 → rank 3), potentially reflecting COVID-19-associated endothelial activation engaging complement pathways.

Mann-Whitney U tests comparing each disease condition to healthy reference showed significant differences for eGFR in DKD (P = 0.031), T2D in DKD (P = 0.018), and IgAN in COV-AKI (P = 0.042). Cell-type rank change analysis across all six traits and five conditions identified 23 instances of rank shifts ≥ 5 positions from healthy reference, concentrated in DKD (9 instances) and AKI (7 instances). Bump plots tracking cell-type enrichment rank changes across conditions for all six traits revealed the magnitude and direction of these shifts, with bold lines highlighting the most dramatic remodeling events (**Fig. 4l–q**).

### Gene-level scDRS correlation identifies condition-specific molecular programs

To identify individual genes whose expressions most strongly track with scDRS disease relevance scores, we computed Spearman rank correlations between gene expression and scDRS norm_score for all expressed genes across each condition-trait combination (**Fig. 5; Extended Data Figs. 7**). In the healthy reference condition, the top correlated genes reflected established biological pathways for each trait. For eGFR, ACSM2B (ρ = 0.34), ACSM2A (ρ = 0.33), and MIOX (ρ = 0.31) topped the list-all encoding enzymes of proximal tubule fatty acid and inositol metabolism^1^. For eGFRcys, the top genes were exclusively mitochondrial-encoded: MT-ATP6 (ρ = 0.31), MT-ND2 (ρ = 0.31), and MT-CYB (ρ = 0.31), reflecting the distinct mitochondrial biology underlying cystatin C-based filtration estimates. BUN’s top genes included KCNJ1 (ρ = 0.18), CDH16 (ρ = 0.17), and CLCNKB (ρ = 0.17), all encoding ion channels and adhesion molecules critical for loop of Henle function. T2D and UACR shared PDE4D as their top correlated gene (T2D: ρ = 0.30; UACR: ρ = 0.40), a phosphodiesterase linked to cAMP signaling and inflammatory pathways^15,16^. For IgAN, the top genes included B2M (ρ = 0.42), TMSB10 (ρ = 0.38), and ribosomal proteins (RPS23, RPS18), consistent with immune activation and antigen presentation pathways.

**Figure 5.**
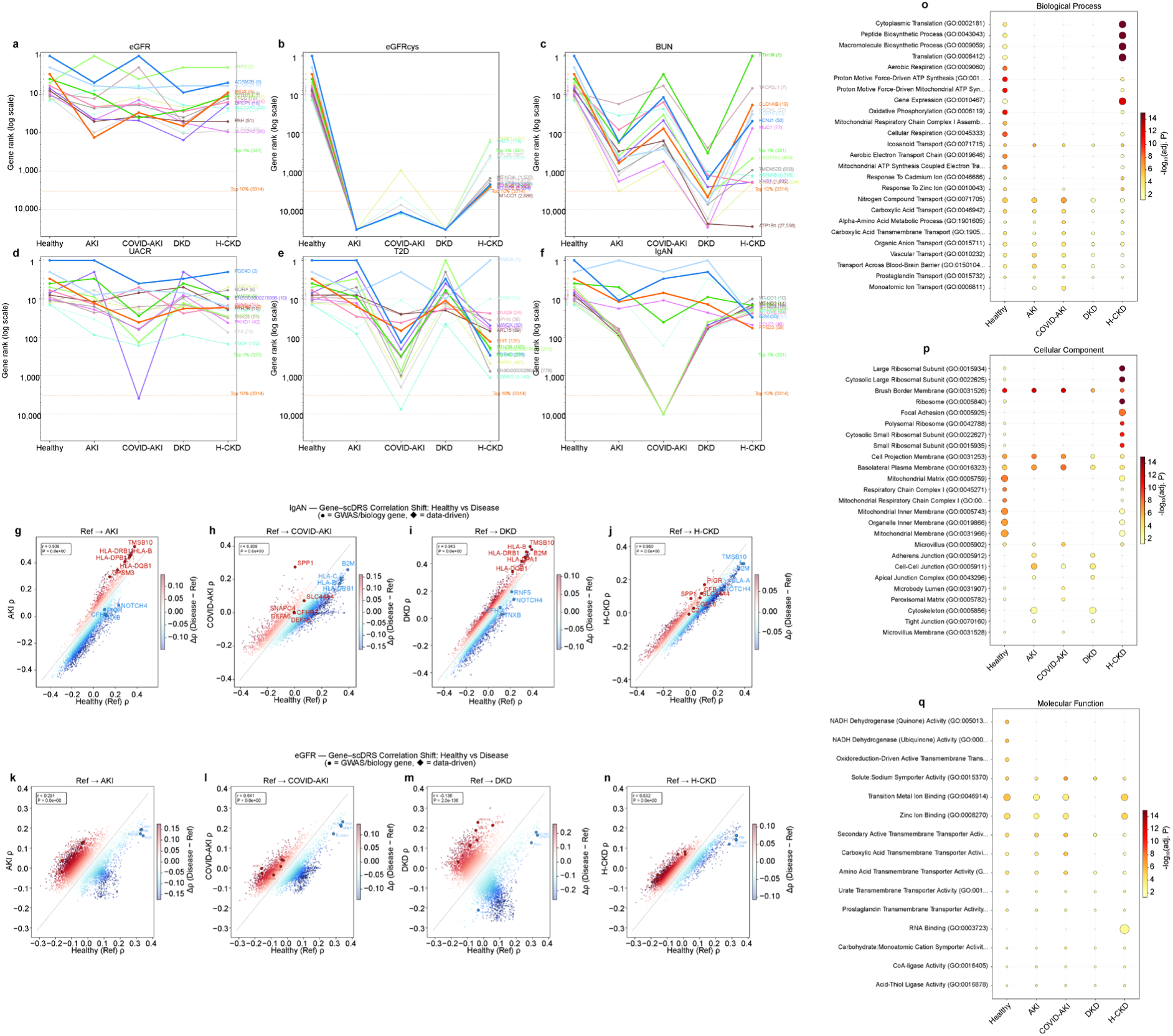
Gene-level correlation analysis and functional pathway characterization. (a–f) Bump charts tracking the rank of the top 15 healthy-reference genes across five disease conditions for each trait: (a) eGFR, (b) eGFRcys, (c) BUN, (d) T2D, (e) UACR, (f) IgAN. Y-axis uses a log scale to capture the full rank range (1 to ∼33,000 genes). Right-side labels show gene name and final rank. Green and orange dashed lines mark the top 1% and top 10% thresholds. (g–j) Scatter plots of Spearman ρ (gene expression vs. scDRS score) in healthy reference (x-axis) versus each disease condition (y-axis) for IgAN: (g) AKI, (h) COV-AKI, (i) DKD, (j) H-CKD. Points colored by Δρ; labeled genes include GWAS-prioritized (circles) and data-driven (diamonds) candidates. (k–n) Same scatter plot format for eGFR across four disease conditions. Pearson r and P-values shown for genome-wide correlation. (o–q) Gene ontology enrichment analysis: (o) compareCluster-style dot plot showing GO enrichment (Biological Process, Cellular Component, Molecular Function) across five conditions for a representative trait. Dot size = gene overlap count; color = −log₁₀(adjusted P-value); × = not significant. (p) Additional GO enrichment panel. (q) GO Biological Process term gain/loss analysis across all six traits and four disease conditions. Horizontal bars show the number of shared, gained (disease-specific), and lost (reference-specific) GO terms. See Extended Data Figs. 6–12 for all traits and comparisons.

Gene rank bump charts revealed dramatic condition-specific reshuffling of the correlation landscape (**Fig. 5a–f**). Using a log-scale visualization to capture the full rank range (1 to ∼33,000 genes), we observed that for eGFR, metabolic genes such as ACSM2B maintained high ranks across conditions (rank 1–5 across all), while transport genes like PAH dropped from rank 8 in healthy to rank 51 in H-CKD. For eGFRcys, mitochondrial genes showed catastrophic rank drops in AKI (MT-ATP6: rank 1 → rank 2,219), consistent with mitochondrial dysfunction during acute injury. BUN genes showed the most extreme condition-dependent behavior: ATP1B1 dropped from rank 8 in healthy to rank 27,558 in H-CKD, and MUC1 fell from rank 10 to rank 29,277 in DKD. These extreme shifts highlight how disease fundamentally rewires the gene-disease relevance landscape.

The Ref-vs-Disease scatter plots with biologically motivated gene labeling (**Fig. 5g–n; Extended Data Fig. 7**) quantified the genome-wide correlation structure shift. For IgAN, the Ref-vs-DKD comparison showed high concordance (Pearson r = 0.84–0.95 across all disease comparisons), consistent with the stable immune-driven enrichment pattern observed at the cell-type level (Fig. 5g–j). For eGFR, the Ref-vs-DKD comparison showed the most dramatic disruption (Pearson r = −0.136, P < 1×10^-4^), indicating near-complete reorganization of the gene-disease correlation landscape in diabetic kidneys (**Fig. 5k–n**). In contrast, eGFR Ref-vs-H-CKD maintained high concordance (r = 0.832), suggesting hypertensive CKD preserves much of the healthy correlation structure.

To characterize the functional programs associated with scDRS-correlated genes, we performed gene ontology (GO) enrichment analysis on the top 500 positively correlated genes per condition-trait combination using three GO libraries: Biological Process, Cellular Component, and Molecular Function (**Fig. 5o–q; Extended Data Fig. 5**). The GO landscape was highly trait- and condition-specific. For eGFR in healthy reference, top enriched terms included organic acid metabolic process, mitochondrial matrix, and oxidoreductase activity—reflecting the metabolic basis of proximal tubular eGFR heritability. In DKD, new terms emerged including extracellular matrix organization, collagen-containing extracellular matrix, and integrin binding, consistent with fibrotic remodeling (**Fig. 5o; Extended Data Fig. 5**). For eGFRcys, the healthy condition was dominated by mitochondrial terms (oxidative phosphorylation, respiratory chain complex, NADH dehydrogenase activity). AKI showed dramatic loss of mitochondrial terms and gain of stress response pathways (**Extended Data Fig. 5**). For BUN, the healthy reference was enriched for ion transport terms (sodium ion transmembrane transport, chloride channel activity), which were largely preserved in H-CKD but disrupted in DKD (**Extended Data Fig. 5**).

T2D GO analysis revealed condition-specific pathway gains: DKD gained terms related to extracellular matrix and cell adhesion, while AKI gained immune and inflammatory response terms (**Extended Data Fig. 5**). IgAN maintained a stable immune-related GO profile across conditions, with terms such as MHC class II protein complex, antigen processing and presentation, and immunoglobulin production consistently enriched (**Extended Data Fig. 5**). UACR showed podocyte-related enrichment in healthy reference (podocyte development, slit diaphragm) that intensified in DKD (**Extended Data Fig. 5**). The GO gain/loss analysis across all six traits and four disease conditions (**Extended Data Fig. 6**) identified DKD as the most disruptive condition, with the highest number of gained and lost GO terms across all traits.

### Disease-specific druggable target nomination

A key translational contribution of our scDRS framework is the ability to nominate druggable targets whose genetic risk relevance emerges specifically in disease contexts. By comparing gene-scDRS correlation ranks between healthy reference and disease conditions, we identified genes with dramatic rank shifts that encode known drug targets or are in active therapeutic development (**Fig. 6a–b**).

**Figure 6.**
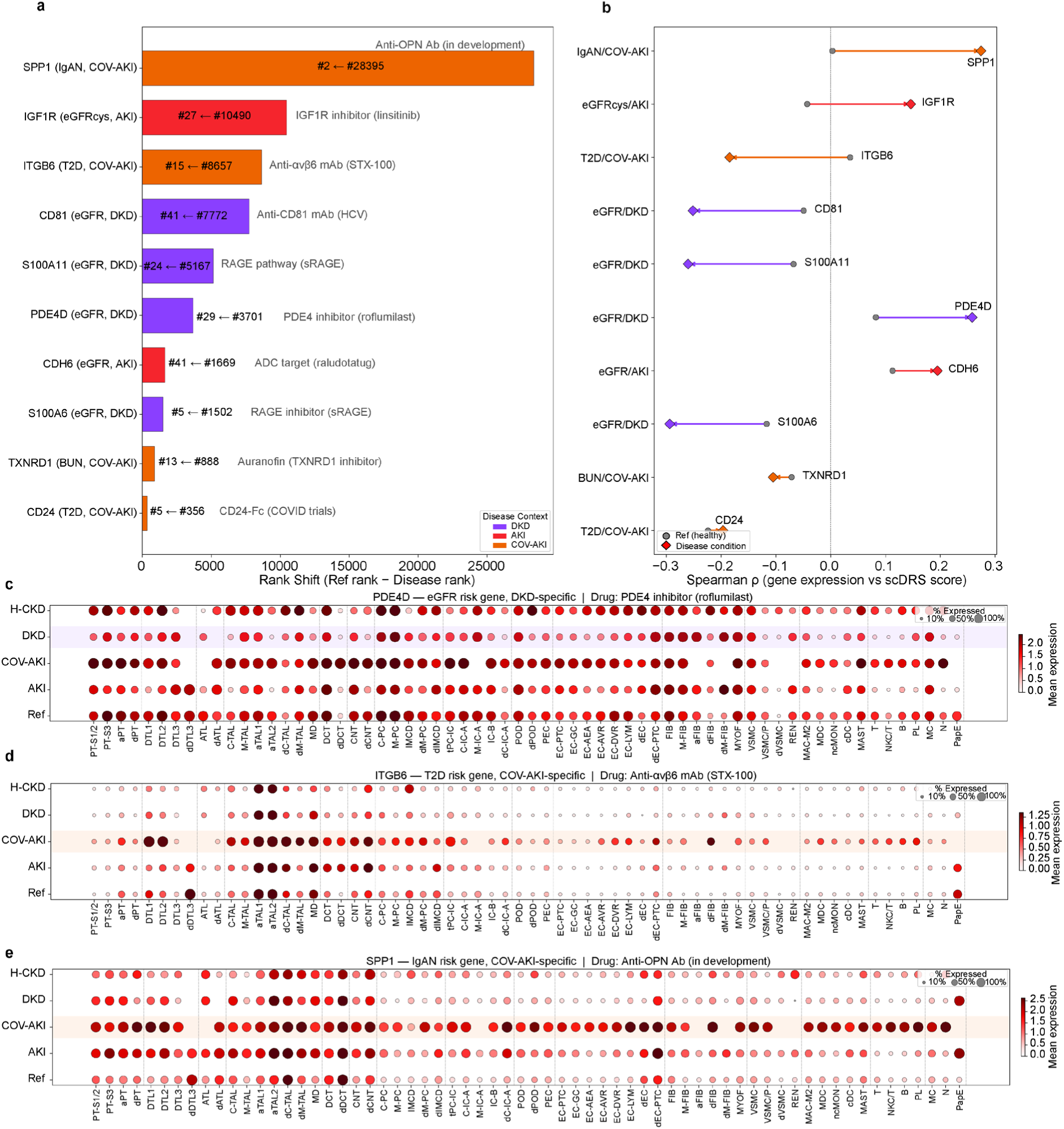
Disease-specific druggable target genes nominated by scDRS. (a) Rank shift waterfall plot for three prioritized druggable targets showing the magnitude of rank change from healthy reference to disease condition. Bar length represents the absolute rank shift; annotations show the reference and disease ranks with corresponding drug candidates. (b) Dumbbell plot showing the Spearman ρ correlation change from healthy reference (circle) to disease condition (diamond) for each target gene. Line length represents the magnitude of correlation shift. (c) Expression dot plot for PDE4D across 73 cell subtypes (L2) and five conditions. Dot size = percentage of cells expressing the gene; color = mean expression level. Disease-specific condition highlighted. PDE4D shows DKD-specific upregulation in PT segments and fibroblasts. Drug: PDE4 inhibitor (roflumilast). (d) Expression dot plot for ITGB6. COV-AKI-specific upregulation in proximal tubule and collecting duct cells. Drug: anti-αvβ6 mAb (STX-100/BG00011). (e) Expression dot plot for SPP1 (osteopontin). COV-AKI-specific broad upregulation across tubular, macrophage, and endothelial populations. Drug: anti-OPN antibody (in development). See Extended Data Fig. 13 for the full set of 10 druggable targets.

We prioritized three high-confidence targets for the main analysis based on rank shift magnitude, biological plausibility, and drug development stage: PDE4D (phosphodiesterase 4D) showed a striking rank shift from position 3,701 in healthy reference to rank 29 for eGFR in DKD, representing a >100-fold improvement in disease relevance. PDE4 inhibitors such as roflumilast are FDA-approved for COPD and have shown renal-protective effects in preclinical models by suppressing inflammation through cAMP-mediated pathways^15,16^. Expression dot plot analysis across 73 cell subtypes revealed that PDE4D upregulation in DKD was concentrated in proximal tubule segments (PT-S1/2, PT-S3) and fibroblasts, suggesting tubular and interstitial targets for PDE4 inhibition (**Fig. 6c**).

ITGB6 (integrin αvβ6) emerged from rank 8,657 in healthy reference to rank 15 for T2D in COV-AKI, reflecting activation of TGF-β signaling and fibrotic pathways during acute injury superimposed on metabolic disease. The anti-αvβ6 monoclonal antibody STX-100 (now BG00011) has been investigated in clinical trials for fibrosis^17^. ITGB6 expression in COV-AKI was concentrated in proximal tubule cells and collecting duct principal cells, with minimal expression in healthy reference (**Fig. 6d**).

SPP1 (osteopontin) showed the most dramatic rank shift of any gene: from rank 28,395 in healthy reference to rank 2 for IgAN in COV-AKI, representing an almost complete switch from irrelevance to top-ranked disease gene. Anti-OPN antibodies are in preclinical development for kidney fibrosis. SPP1 expression analysis revealed broad upregulation in COV-AKI across proximal tubule, macrophage, and endothelial populations, consistent with its role as a secreted cytokine in kidney inflammation and injury repair (**Fig. 6e**).

Seven additional druggable targets were identified with notable rank shifts, including IGF1R (linsitinib; eGFRcys/AKI), CD81 (anti-CD81 mAb; eGFR/DKD), S100A11/S100A6 (RAGE pathway; eGFR/DKD), CDH6 (raludotatug ADC; eGFR/AKI), TXNRD1 (auranofin; BUN/COV-AKI), and CD24 (CD24-Fc; T2D/COV-AKI). Full rank shift analysis and expression dot plots for all ten candidates are provided in **Extended Data Fig. 9**.

## Discussion

We present the Kidney Genetic Disease Cell Atlas, a comprehensive single-cell resolution map of genetic risk for six kidney-related traits across five clinical conditions. By integrating GWAS summary statistics with single-cell and spatial transcriptomics through the scDRS framework, we provide the most detailed characterization to date of how kidney disease heritability is distributed across cell types and how this distribution is remodeled by disease.

The dominance of proximal tubule cells for eGFR heritability is consistent with prior findings from Sheng et al.^13^ using stratified LD score regression and from Muto et al.^14^ using snRNA-seq-based approaches. However, our analysis extends these findings in several important ways. First, by applying scDRS rather than aggregate enrichment methods, we obtain cell-level resolution that reveals heterogeneity within cell types. For example, not all PT cells are equally enriched for eGFR heritability, with PT-S1/2 segments showing stronger signals than PT-S3. Second, our condition-specific analysis reveals that the PT dominance for eGFR, while stable, is accompanied by disease-specific gains in fibroblasts (DKD) and immune cells (AKI) that would be invisible in healthy-only analyses. Third, the independent spatial validation using Slide-seqV2 data provides orthogonal evidence that our enrichment patterns reflect genuine biological signals rather than technical artifacts of nuclear isolation.

The strong IgAN enrichment in immune cells (Cohen’s d = 1.40) validates the autoimmune basis of this disease and highlights the power of scDRS to capture disease-relevant cell populations. The stability of IgAN immune enrichment across all five conditions (d > 1.0 in all) is remarkable and suggests that the genetic architecture of IgAN susceptibility is intrinsically tied to immune cell biology regardless of kidney health status. This is consistent with the known role of mucosal IgA production, complement activation, and galactose-deficient IgA1 in IgAN pathogenesis^5^. The observation that endothelial cells gain IgAN enrichment specifically in COV-AKI may reflect COVID-19-associated endothelial activation engaging complement pathways that overlap with IgAN susceptibility loci.

The eGFRcys results provide a striking contrast to eGFR, with mitochondrial genes dominating the healthy correlation landscape. This difference likely reflects the distinct genetic architectures of creatinine-based versus cystatin C-based GFR estimation^2^. Cystatin C is freely filtered at the glomerulus and completely reabsorbed and catabolized in proximal tubules, making its levels more dependent on mitochondrial function and tubular health than creatinine clearance. The dramatic loss of mitochondrial gene correlations in AKI (MT-ATP6 dropping from rank 1 to rank 2,219) provides a molecular signature of acute mitochondrial dysfunction during kidney injury.

Our gene-level scDRS correlation analysis complements the cell-type-level analysis by identifying specific genes whose expression most strongly tracks disease relevance. The shared PDE4D dominance between T2D and UACR traits suggests overlapping cAMP-mediated pathways connecting metabolic disease and albuminuria, consistent with epidemiological evidence linking diabetes to proteinuria through both hemodynamic and direct tubular mechanisms^15^. The observation that PDE4D rank shifts from ∼3,700 in healthy to rank 29 in DKD suggests that the diabetic kidney microenvironment specifically amplifies PDE4D-mediated signaling.

The GO enrichment analysis reveals that disease conditions not only shift which genes are most relevant but also remodel the functional pathway landscape. The emergence of extracellular matrix and integrin-related terms for eGFR in DKD, absent in healthy reference, provides pathway-level evidence for fibrotic remodeling engaging GFR-associated genetic programs. The loss of mitochondrial terms for eGFRcys in AKI at the pathway level mirrors the gene-level findings and highlights oxidative phosphorylation as a key pathway disrupted during acute injury.

A key translational contribution is the nomination of disease-specific druggable targets through scDRS rank shift analysis. PDE4D, ITGB6, and SPP1 each represent distinct therapeutic opportunities: PDE4D targets the cAMP/inflammatory axis with an existing FDA-approved inhibitor (roflumilast) that could be repurposed for DKD^16^; ITGB6 targets TGF-β-mediated fibrosis with a clinical-stage antibody (STX-100/BG00011)^17^; and SPP1 targets injury-associated inflammation through anti-osteopontin approaches currently in preclinical development. The cell-type specificity revealed by expression dot plots (**Fig. 6c–e**) provides additional precision for targeting strategies.

Several limitations should be noted. First, scDRS scores depend on the input gene set derived from MAGMA, and traits with smaller GWAS sample sizes (IgAN, n = 10,146 cases) may have less well-powered gene sets. Second, the Lake et al. atlas, while comprehensive, has unequal representation across conditions (COV-AKI: 19,575 cells vs. Ref: 107,701 cells), which may affect statistical power for disease-specific analyses. Third, our spatial validation used healthy tissue only; spatial data from diseased kidneys would strengthen disease-specific claims. Fourth, scDRS captures linear expression-disease correlations and may miss non-linear or interaction effects. Fifth, the druggable target nominations are based on expression correlation patterns and require experimental validation in disease models.

Future directions include applying this framework to additional kidney diseases (FSGS, lupus nephritis, ANCA vasculitis), integrating multi-omic data (ATAC-seq, methylation) to capture regulatory mechanisms, and experimentally validating the nominated drug targets in kidney organoid and animal models^18^. The increasing availability of spatial multi-omics technologies will enable more refined mapping of disease-specific heritability to anatomical niches within the kidney.

In conclusion, the Kidney Genetic Disease Cell Atlas provides a resource for the nephrology and genetics communities to explore how genetic risk for kidney traits maps onto specific cell types and how disease remodels this landscape. The atlas is publicly available and can serve as a foundation for future functional genomics studies and therapeutic development in kidney disease.

## Methods

### GWAS summary statistics

GWAS summary statistics were obtained from published sources: eGFR and BUN from Wuttke et al. (2019)^1^; eGFRcys from Stanzick et al. (2021)^2^; T2D from Mahajan et al. (2022)^4^; UACR from Teumer et al. (2019)^3^; and IgAN from Kiryluk et al. (2023)^5^. All summary statistics were based on European-ancestry populations and were formatted to include columns for SNP identifier, chromosome, base position, effect allele, non-effect allele, Z-score or beta, standard error, and P-value.

### MAGMA gene-level analysis

MAGMA v1.10^11^ was used to compute gene-level association statistics from GWAS summary data. Gene boundaries were defined using NCBI GRCh37/hg19 gene locations, with a 10-kb upstream window to capture promoter-proximal regulatory variants. The 1000 Genomes Phase 3 European reference panel was used for LD estimation. The SNP-wise mean model was applied to aggregate SNP associations within each gene. The top 1,000 genes ranked by MAGMA P-value for each trait were selected as input gene sets for scDRS.

### Single-cell RNA sequencing data

Single-nucleus RNA-seq data from 304,652 cells (37 adult kidney samples, 5 conditions) were obtained from Lake et al. (2023)^6^. Cell type annotations were used at two resolution levels: subclass L1 (16 major types: PT, DTL, ATL, TAL, DCT, CNT, PC, IC, POD, PEC, PapE, EC, VSM/P, FIB, NEU, IMM) and subclass L2 (73 refined subtypes). Condition labels comprised Ref (healthy reference; n = 107,701), AKI (acute kidney injury; n = 65,171), COV-AKI (COVID-19-associated AKI; n = 19,575), DKD (diabetic kidney disease; n = 67,628), and H-CKD (hypertensive CKD; n = 44,577). The h5ad object was processed using Scanpy v1.9^19^.

### Slide-seqV2 spatial transcriptomics

Slide-seqV2 spatial transcriptomics data comprising 920,088 beads from 44 pucks (22 cortex, 22 medulla) of healthy adult kidney tissue from 11 donors were obtained from Marshall et al. (2022)^7^ via GEO accession GSE190094. Cell type annotations were transferred from the snRNA-seq reference using RCTD (Robust Cell Type Decomposition). For spatial visualization, one representative cortex and one medulla puck were selected based on largest bead count. Spatial heatmaps of scDRS scores were generated by pooling results across all 44 pucks for statistical comparisons.

### scDRS computation

scDRS v1.0.2^8^ was used to compute single-cell disease relevance scores. For each trait, the top 1,000 MAGMA-ranked genes were provided as the gene set. scDRS was run using default parameters: 1,000 control gene sets matched by mean expression and expression variance were used for normalization. The output included a normalized disease relevance score (norm_score) and associated P-value for each cell. Cells with P < 0.05 were classified as significantly enriched. For condition-specific analysis, scDRS was applied separately to cells from each clinical condition, using the same MAGMA-derived gene sets.

### Gene-scDRS correlation analysis

For each condition-trait combination, Spearman rank correlations were computed between the expression of each gene and the scDRS norm_score across all cells in that condition. Genes were ranked by absolute correlation magnitude. The top 500 positively and top 500 negatively correlated genes were retained for downstream analysis. Cross-condition rank shifts were calculated as the difference in rank between the healthy reference and each disease condition for each gene. Priority genes for visualization labeling were selected from the intersection of MAGMA top genes and known kidney biology markers.

### Gene ontology enrichment

GO enrichment was performed using gseapy v1.1 with the Enrichr backend. Three GO libraries were queried: GO_Biological_Process_2023, GO_Cellular_Component_2023, and GO_Molecular_Function_2023. The top 500 positively correlated genes per condition-trait combination were used as input. Terms with Benjamini-Hochberg adjusted P < 0.05 were considered significant. Cross-condition comparison identified GO terms gained (significant in disease but not reference) and lost (significant in reference but not disease) for each trait-disease pair.

### Druggable target nomination

Druggable targets were identified by intersecting genes with large rank shifts (Δrank > 500 between healthy reference and disease) with the DGIdb druggable genome database and manual literature curation. For each candidate, we verified (1) the existence of a drug or drug candidate targeting the gene product, (2) the biological plausibility of the gene’s role in kidney disease, and (3) the cell-type specificity of expression changes using dot plots across 73 L2 subtypes and 5 conditions.

### Statistical analysis

All statistical analyses were performed in Python 3.10 using scipy v1.11, numpy v1.24, and pandas v2.0. Cell-type enrichment significance was assessed using Wilcoxon rank-sum tests comparing scDRS scores of each cell type against all other cells, with Bonferroni correction for multiple comparisons. Condition-specific comparisons used Mann-Whitney U tests. Cohen’s d effect sizes were computed as the difference in mean scDRS scores divided by the pooled standard deviation. Spearman rank correlations were used for cross-platform comparisons and gene-level analyses. All figures were generated using matplotlib v3.7 at 600 DPI with PDF font type 42 for vector text editability. Visualization code is available in the accompanying GitHub repository.

---

**Extended Data Figure 1.**
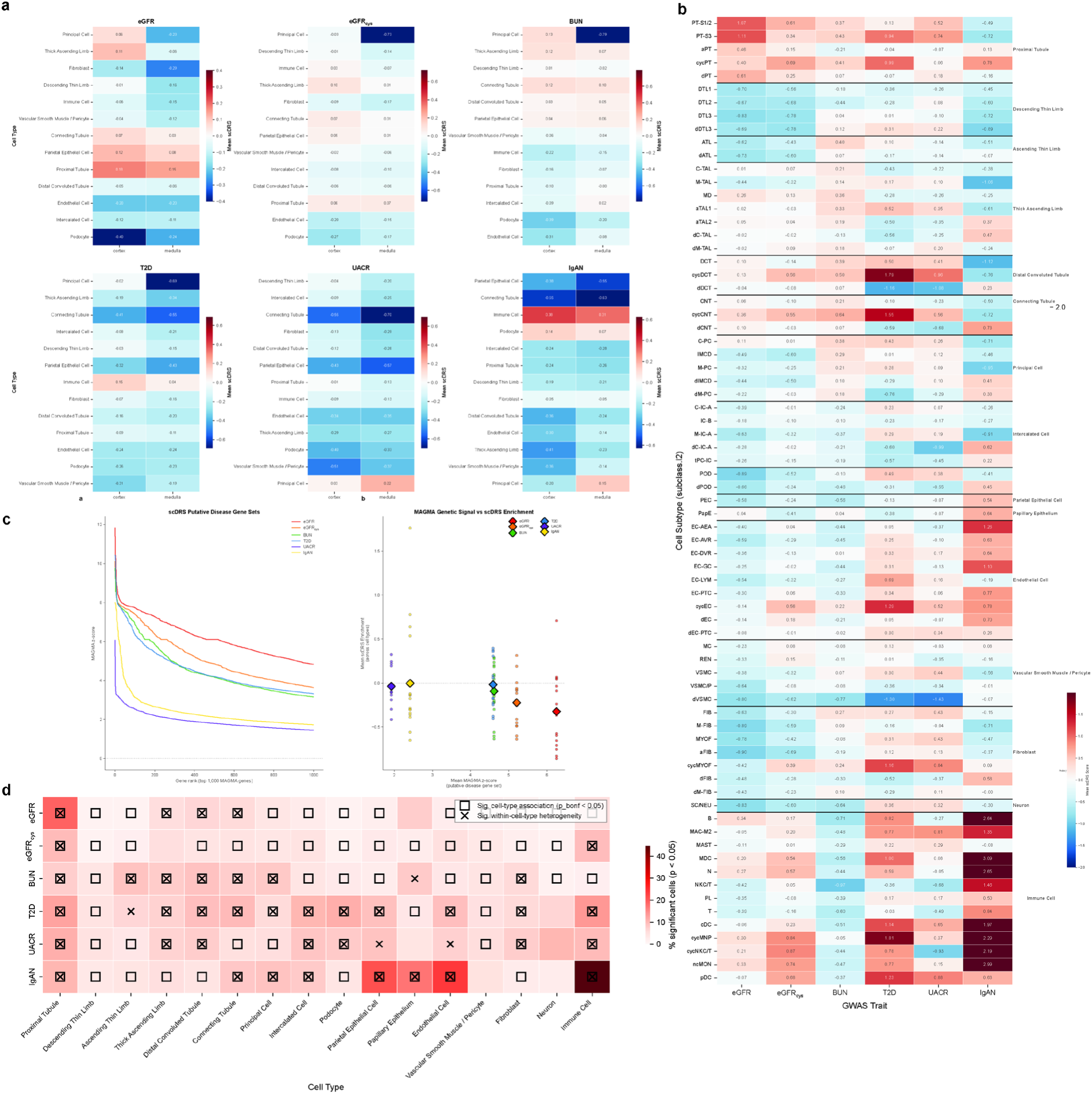
Detailed cell-type enrichment analysis. (a) Heatmap of mean scDRS scores across cell types and traits, stratified by cortex/medulla region. (b) Heatmap at the L2 subtype level (73 subtypes × 6 traits) showing fine-grained enrichment patterns. (c) MAGMA vs. scDRS comparison: scatter plot of gene-level MAGMA Z-scores versus scDRS-derived cell-type enrichment effect sizes. (d) Proportion of significant cells (P < 0.05) per cell type for each trait in healthy reference.

**Extended Data Figure 2.**
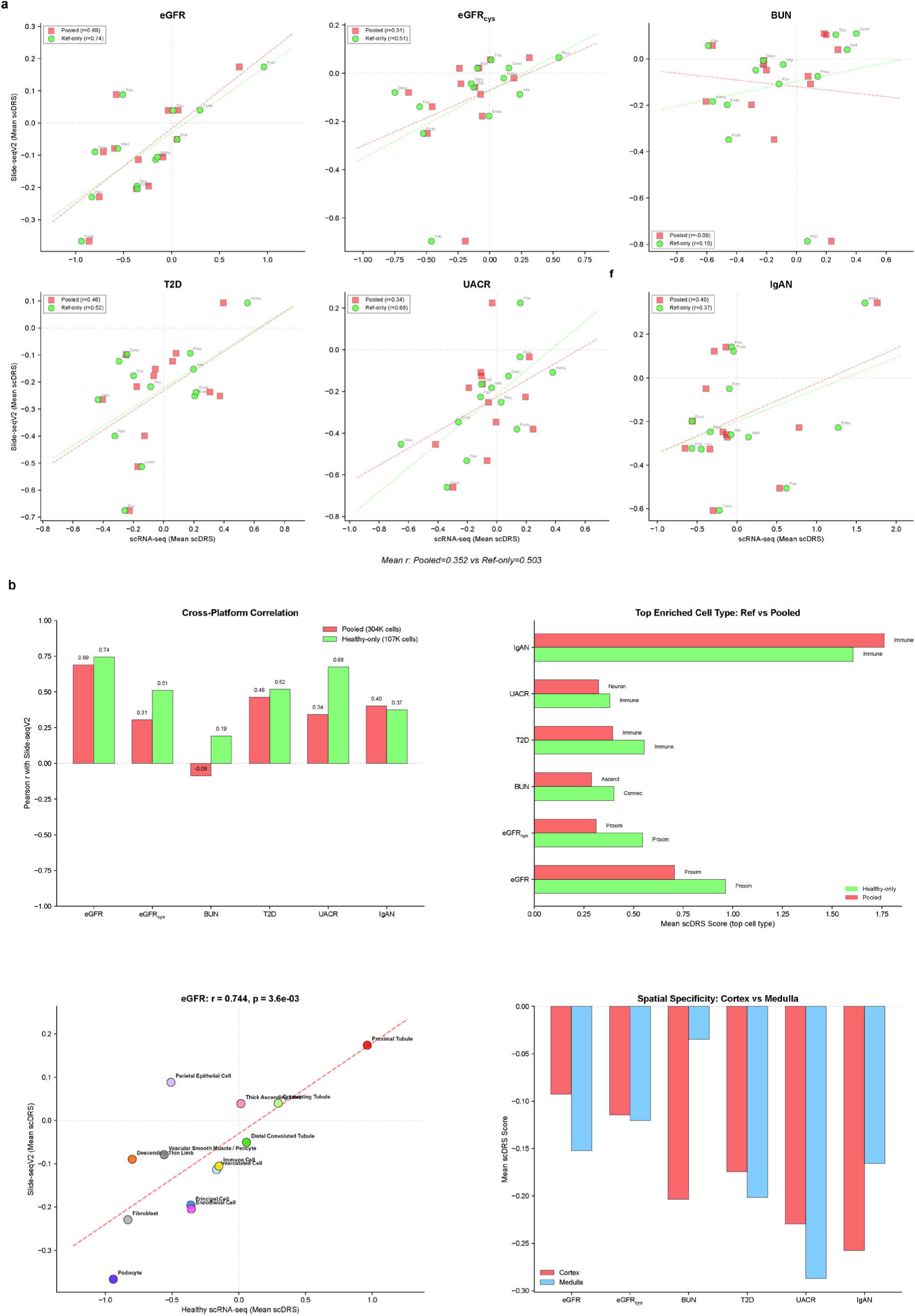
Spatial transcriptomics validation details. (a) Comparison of pooled (all pucks) vs. reference-only Slide-seqV2 enrichment. (b) Comprehensive healthy tissue validation panel including per-puck variability and cross-platform agreement metrics.

**Extended Data Figure 3.**
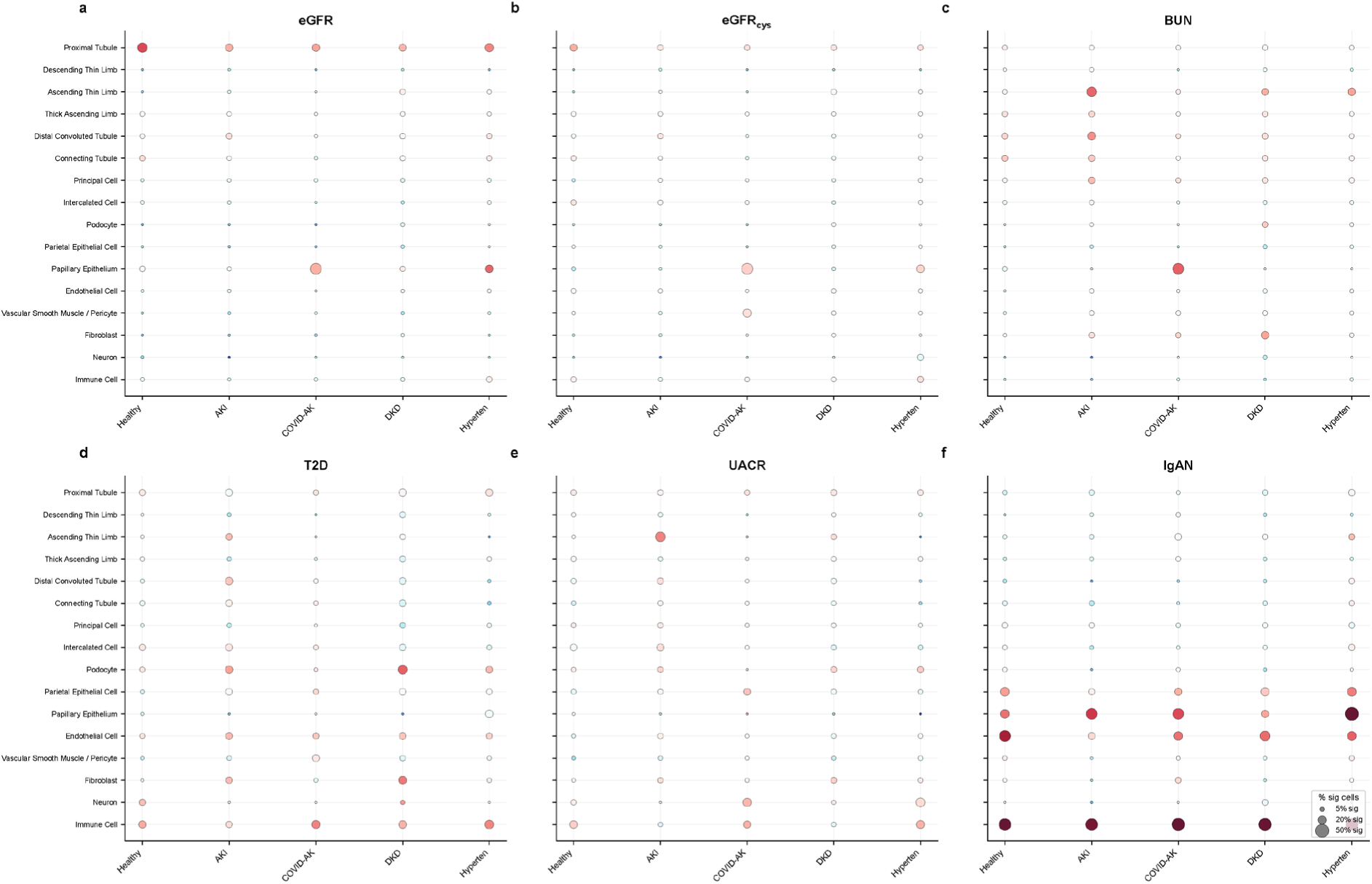
Disease-condition-specific analysis details. Bubble plots showing condition-specific enrichment changes with size proportional to cell count and color reflecting direction of change.

**Extended Data Figure 4.**
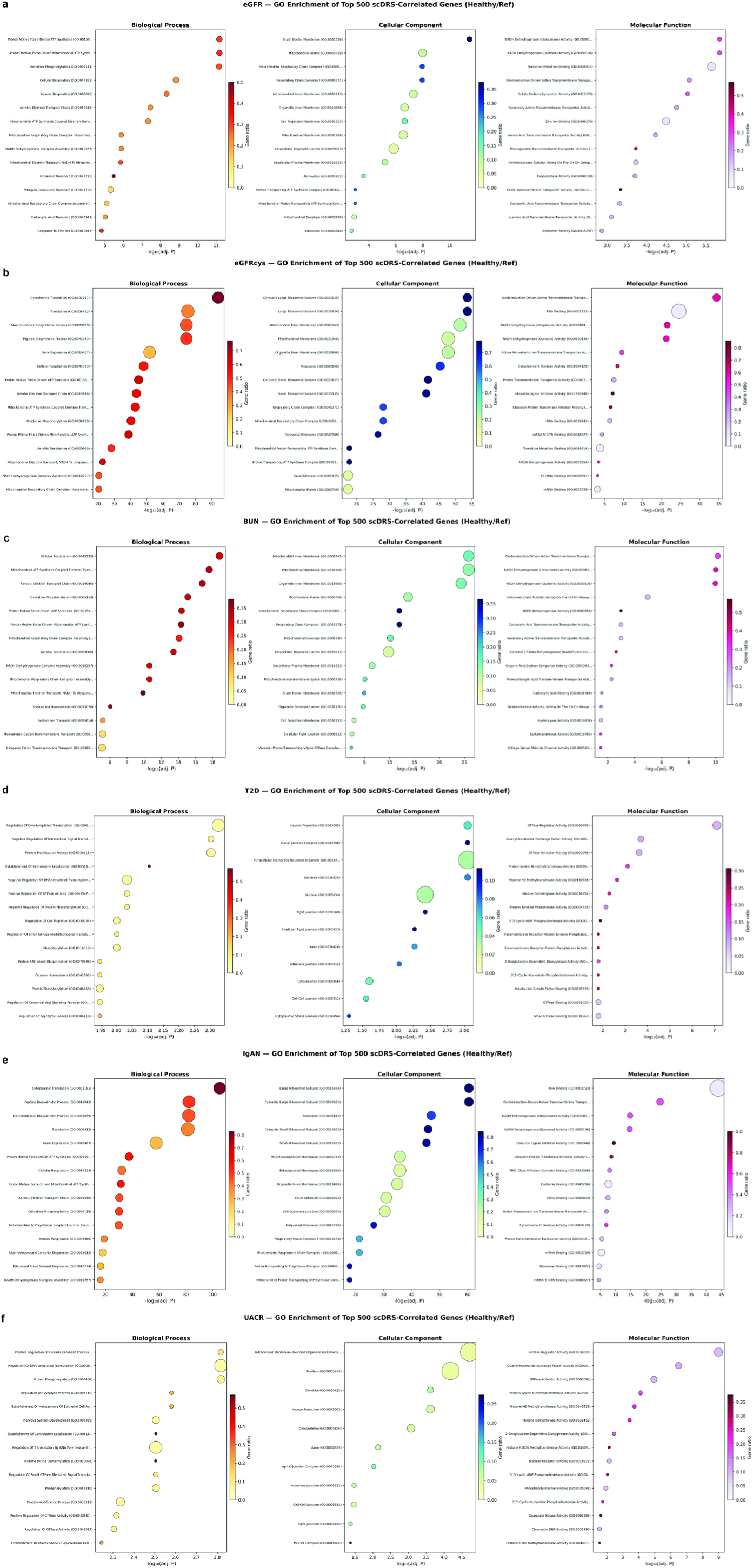
Gene ontology enrichment dot plots. (a–f) GO enrichment dot plots for each trait: (a) eGFR, (b) eGFRcys, (c) BUN, (d) T2D, (e) IgAN, (f) UACR. Each panel shows the top enriched terms from Biological Process, Cellular Component, and Molecular Function across all five conditions.

**Extended Data Figure 5.**
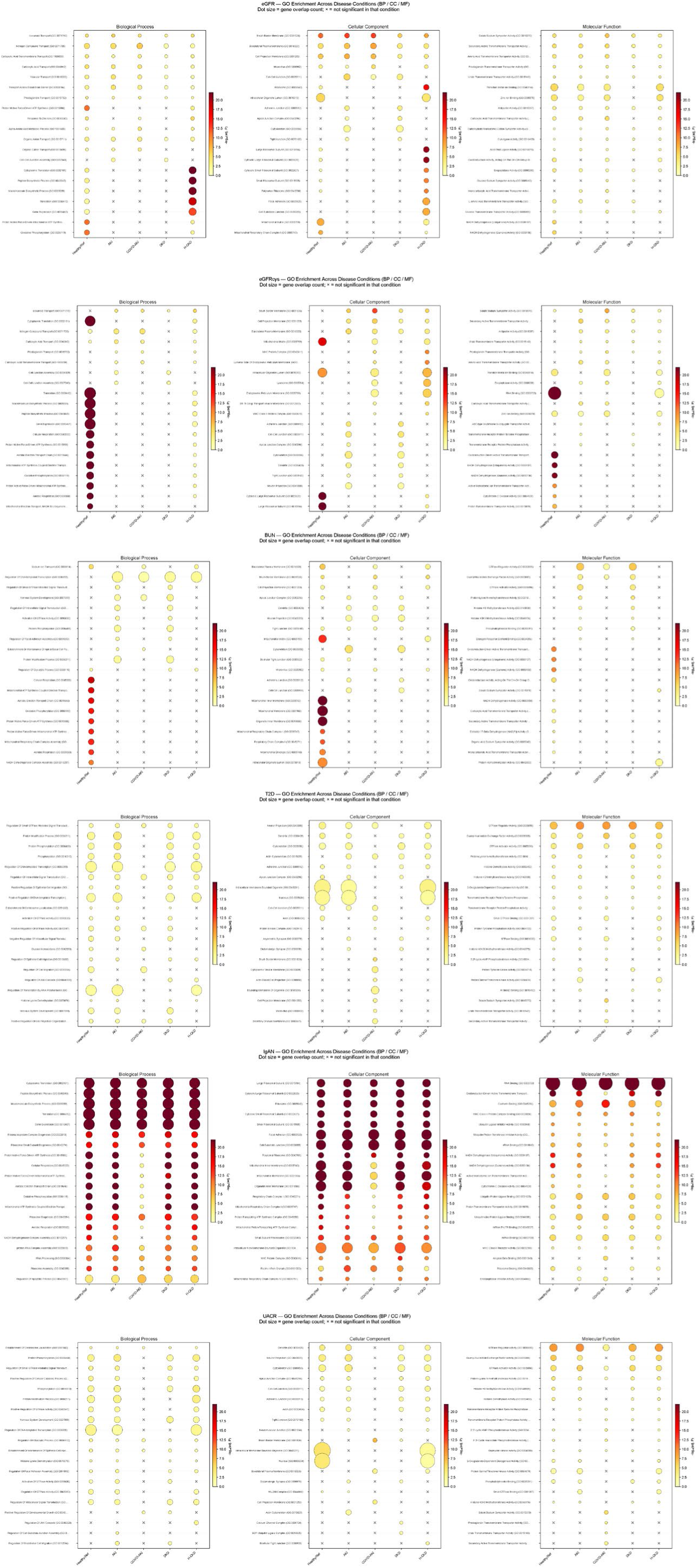
Cross-condition GO term comparison. (a–f) Heatmaps comparing GO term significance across conditions for each trait: (a) eGFR, (b) eGFRcys, (c) BUN, (d) T2D, (e) IgAN, (f) UACR. Color reflects −log₁₀(adjusted P); rows are GO terms; columns are conditions.

**Extended Data Figure 6.**
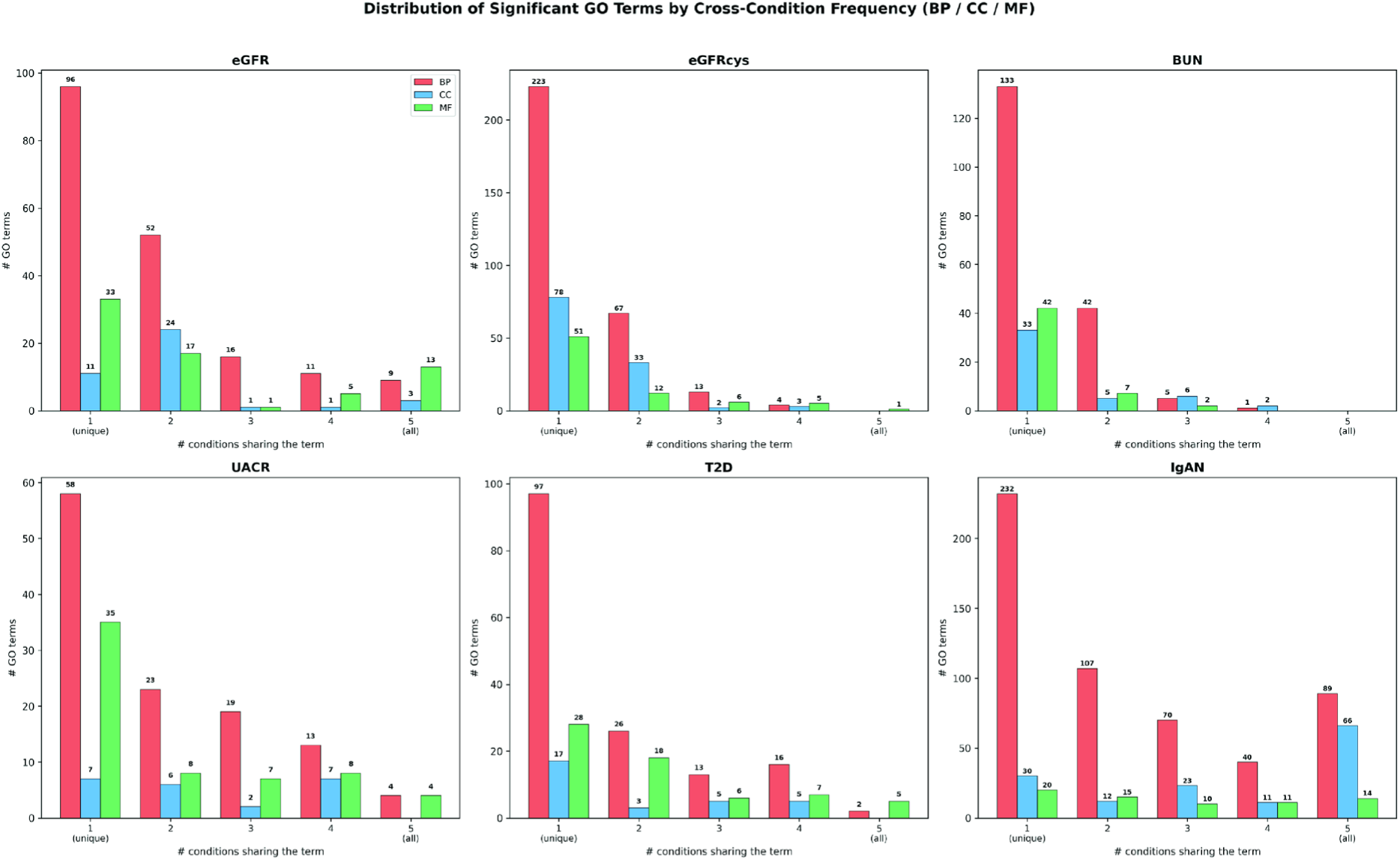
GO term shared vs. condition-specific analysis. Summary showing the number of shared, condition-specific gained, and condition-specific lost GO Biological Process terms for each trait-disease comparison across all six traits and four disease conditions.

**Extended Data Figure 7.**
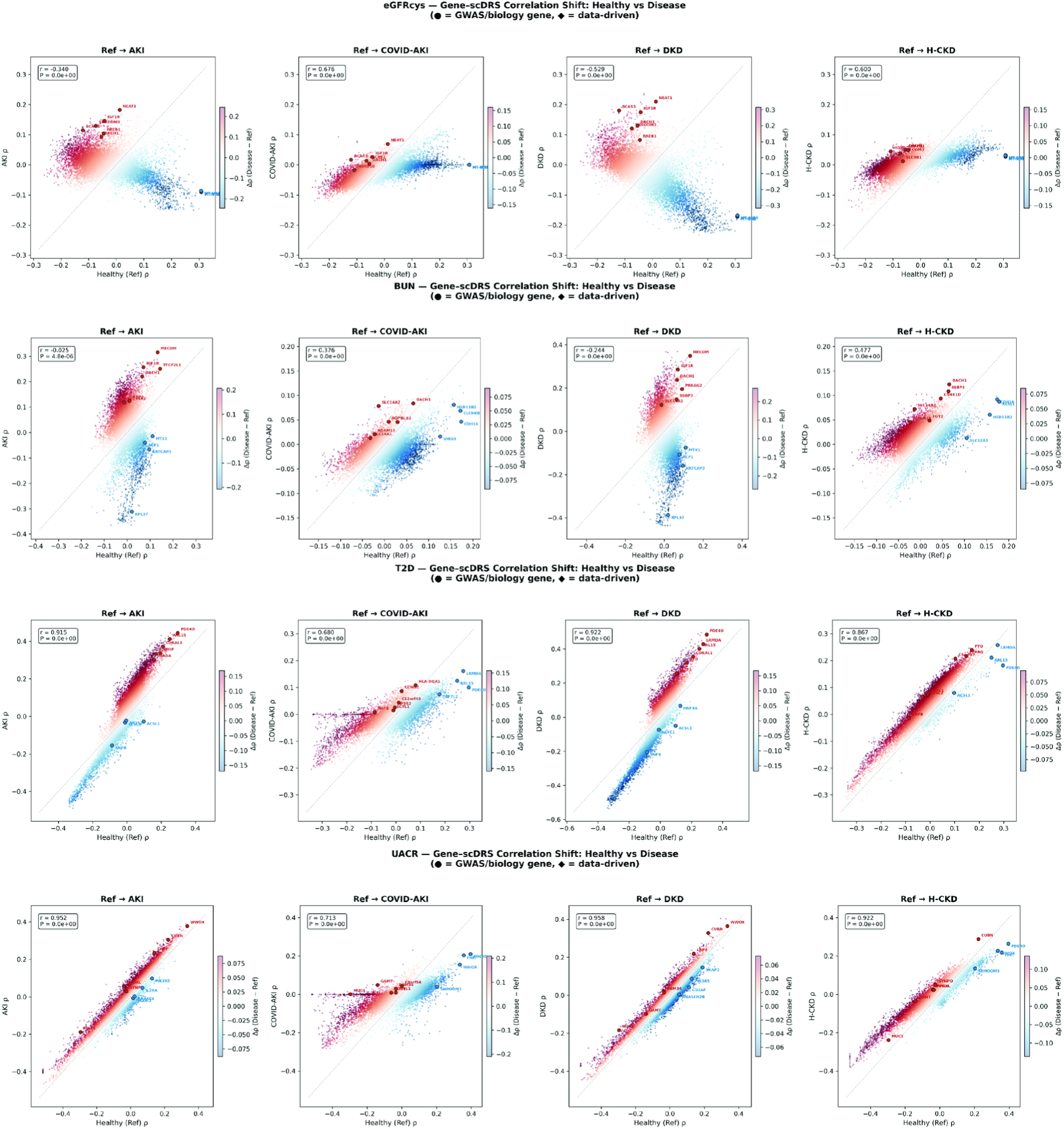
Gene-scDRS correlation scatter plots for additional traits. eGFRcys: Ref ρ vs. disease ρ scatter plots for all four disease conditions with labeled priority genes. BUN. T2D. UACR.

**Extended Data Figure 8.**
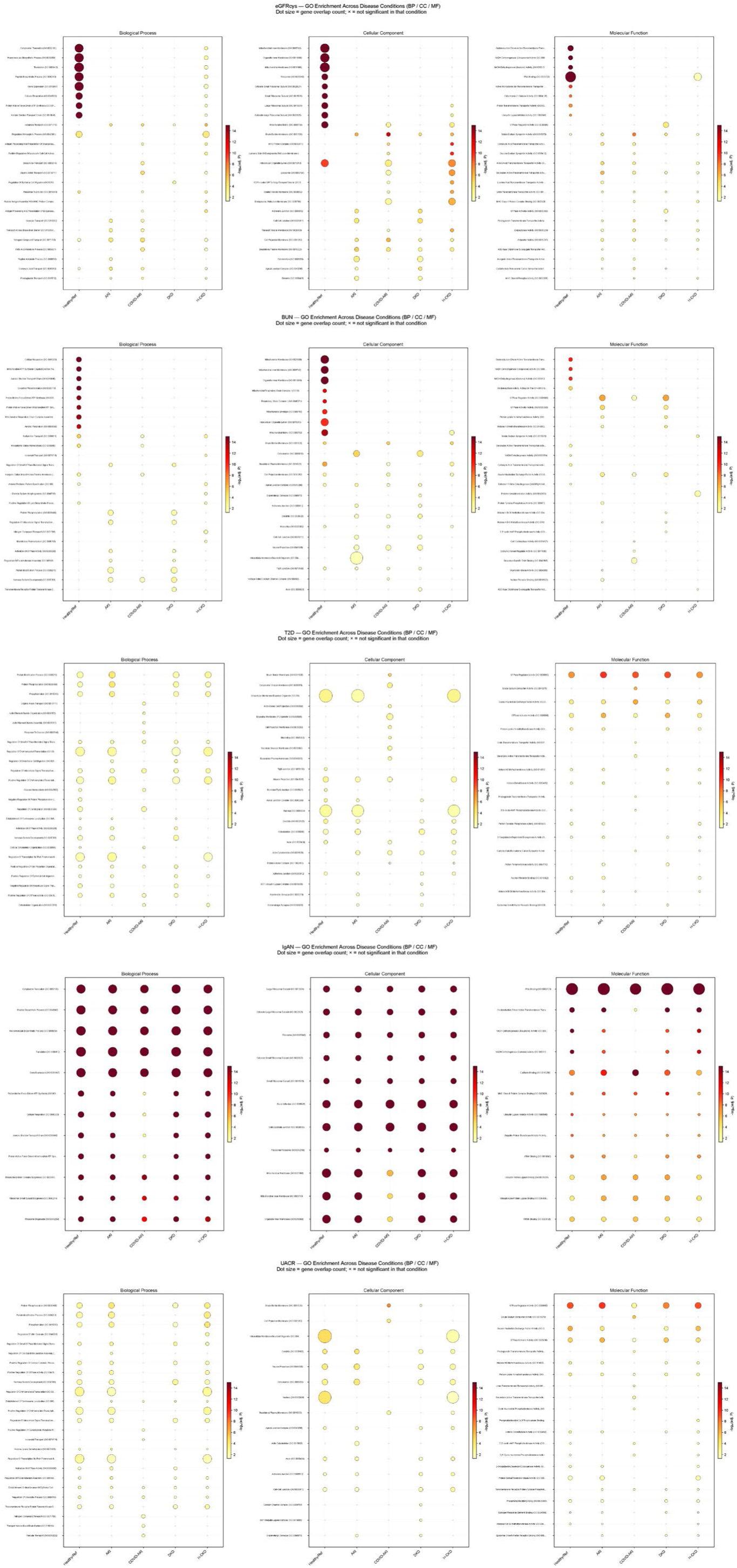
compareCluster GO enrichment dot plots for additional traits. eGFRcys, BUN, T2D, IgAN, UACR.

**Extended Data Figure 9.**
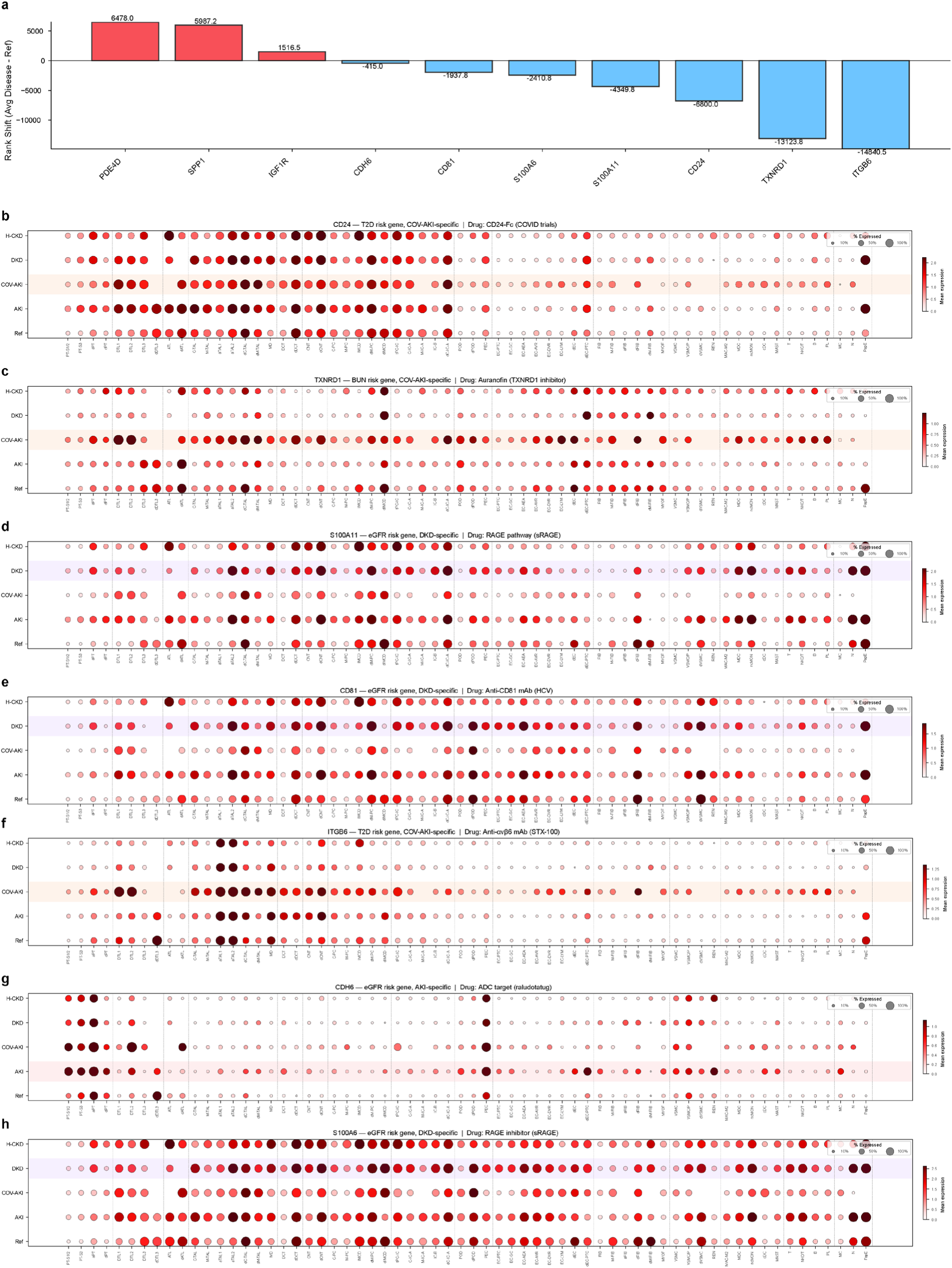
Extended druggable target analysis. (a) Rank shift bar chart for all 10 druggable candidate genes. (b–h) Expression dot plots across 73 cell subtypes and 5 conditions for the 7 additional druggable targets not shown in the main figure: IGF1R (linsitinib), CD81 (anti-CD81 mAb), S100A11 (RAGE pathway/sRAGE), CDH6 (raludotatug ADC), S100A6 (RAGE inhibitor/sRAGE), TXNRD1 (auranofin), CD24 (CD24-Fc). Each dot plot shows condition-specific expression changes with the disease of maximum rank shift highlighted.

